# Niclosamide reverses SARS-CoV-2 control of lipophagy

**DOI:** 10.1101/2021.07.11.451951

**Authors:** Timothy J. Garrett, Heather Coatsworth, Iqbal Mahmud, Timothy Hamerly, Caroline J. Stephenson, Hoda S. Yazd, Jasmine Ayers, Megan R. Miller, John A. Lednicky, Rhoel R. Dinglasan

## Abstract

The global effort to combat COVID-19 rapidly produced a shortlist of approved drugs with anti-viral activities for clinical repurposing. However, the jump to clinical testing was lethal in some cases as a full understanding of the mechanism of antiviral activity as opposed to pleiotropic activity/toxicity for these drugs was lacking. Through parallel lipidomic and transcriptomic analyses we observed massive reorganization of lipid profiles of infected Vero E6 cells, especially plasmalogens that correlated with increased levels of virus replication. Niclosamide (NIC), a poorly soluble anti-helminth drug identified for repurposed treatment of COVID-19, reduced the total lipid profile that would otherwise amplify during virus infection. NIC treatment reduced the abundance of plasmalogens, diacylglycerides, and ceramides, which are required for virus production. Future screens of approved drugs may identify more druggable compounds than NIC that can safely but effectively counter SARS-CoV-2 subversion of lipid metabolism thereby reducing virus replication. However, these data support the consideration of niclosamide as a potential COVID-19 therapeutic given its modulation of lipophagy leading to the reduction of virus egress and the subsequent regulation of key lipid mediators of pathological inflammation.

## INTRODUCTION

The current pandemic spread of SARS-CoV-2 has resulted in >600,000 deaths in the United States alone and >176M cases and >3.8M deaths worldwide (Dong et al., 2020). In 2020, there was a rapid global effort to study severe acute respiratory syndrome coronavirus 2 (SARS-CoV-2) infection kinetics *in vitro* to identify pathways that are involved in entry, replication, and egress of the virus as new targets that can be treated with existing drugs (Gautret et al., 2020; Jeon et al., 2020; Liu et al., 2020; Vincent et al., 2005; Wang et al., 2020; Zhou et al., 2020). However, this worldwide effort was fraught with inconsistencies ranging from the mammalian host cell used for infection studies, to overall study design, which made it difficult to identify common host cell pathways critical to virus biology that can be targeted with available compounds.

Chloroquine (CQ) and, hydroxychloroquine (HCQ), which are indicated for treating malaria, have been shown to also have antiviral activity against SARS-CoV-2 (D’Alessandro et al., 2020; Dyall et al., 2014; Liu et al., 2020; Mauthe et al., 2018; Vincent et al., 2005; Wang et al., 2020). However, the direct and rapid translation of this finding to clinical studies during the pandemic was either equivocal at best, or disastrous (Axfors et al., 2021). Although no studies to date have specifically and comprehensively explored the underlying antiviral mechanism of action (MOA) for these two drugs, they nonetheless moved quickly to COVID-19 clinical studies. CQ and HCQ are 4-aminoquinolines (4-AQ), and another human-safe 4-AQ drug, Amodiaquine (AQ), has also been shown to have antiviral activity against SARS-CoV-1 and Middle East respiratory syndrome coronavirus (MERS-CoV) (Dyall et al., 2014; Vincent et al., 2005), as well as Ebola (D’Alessandro et al., 2020), suggesting that there are common MOAs underpinning broad 4-AQ antiviral activity. Initially, these 4-AQ compounds were considered as candidate partner drugs that can be used along with remdesivir (Wang et al., 2020), the leading broad-spectrum RNA synthesis inhibitor that has shown some clinical benefit (Goldman et al., 2021). The exact nature of the cellular pathways and processes that are targeted by these 4-AQs resulting in reductions in virus propagation remain unknown, but direct protein interaction studies with CQ suggest broad pleiotropic effects on the cell (Gordon et al., 2020). The pleiotropic effects of drugs such as CQ should have given pause, preventing its quick translation to the clinic wherein higher doses of this drug exacerbate the pleiotropic side effects that can result in direct harm to the subject in clinical trials (Borba et al., 2020).

Another anti-parasitic drug, niclosamide (NIC), which suppresses MERS-CoV propagation by inhibiting autophagosome-lysosome fusion through the Beclin1 (Bec1, autophagy regulator/antiapoptotic protein)-SKP2 (S-phase kinase-associated protein 2) pathway (Gassen et al., 2019) has also been proposed to be a potent candidate drug for repurposing in the treatment of SARS-CoV-2 (Xu et al., 2020). NIC is a salicylanilide, oral anti-helminthic drug used in human and veterinary medicine. Akin to the 4-AQs, it appears to have functional effects beyond the Bec1-SKP2 axis in a cell, resulting in broad anti-infective properties, and as a result, also suffers from poor translational potential to the clinic for COVID-19. Considering the failure of CQ and HCQ to progress towards clinical utility, the candidacy of NIC, which is at an earlier stage of consideration for COVID-19 treatment, compelled us to examine more closely its global effects on host cells to determine if we can uncouple the reported antiviral MOA via the Bec1-SKP2 pathway as well as other putative antiviral mechanisms from intrinsic cytotoxicity. We hypothesize that NIC exerts pleiotropic functional activities on a host cell that result in the perturbation of two intimately associated cellular pathways: autophagy and lipid metabolism. These two processes are dysregulated at critical SARS-CoV-2 life cycle checkpoints during virus infection (Stukalov et al., 2021), and treatment with NIC is hypothesized to reverse this dysregulation. Recently, a multi-omics analysis of clinical samples from COVID-19 patients also described a marked change in lipidomics profiles that correlated with disease severity (Overmyer et al., 2021), but this approach could not characterize the mechanism at the cellular level. Herein, we identified and characterized autophagic or lipophagic pathways (and wider linked cellular networks) that are targeted by NIC in the presence and absence of a SARS-CoV-2 infection. We used the lipid signatures induced by this anti-parasitic compound to detect pathways that are predicted to either lead to direct antiviral effects or to cellular dysregulation and cell death. We discuss the utility of this approach in informing the selection of repurposed drugs that may have more clinical benefit, better bioavailability, and lesser cytotoxicity in the context of COVID-19 and other infectious diseases.

## RESULTS

### SARS-CoV-2 infection alters the host cell lipidome

The mechanism by which SARS-CoV-2 enters host cells and systematically alters the cellular environment is currently one of the most compelling areas of study to support the development of antiviral interventions (V’kovski et al., 2020). Evidence suggests that SARS-CoV-2 subverts pre-existing cellular lipids and lipid signaling mechanisms for entry, intracellular trafficking and egress (Abu-Farha et al., 2020). This is consistent with what is known about viruses in general (Mazzon and Mercer, 2014). To investigate the role of lipids in SARS-CoV-2 during the infection cycle in host cells, we profiled the global lipidome from Vero E6 cells at 16h and 48h post-infection with SARS-CoV-2 (**Figure S1A**). The correlation matrix across the groups studied showed a distinct clustering in SARS-CoV-2 infected cells as well as clear clustering with virus and NIC treatment (**Figure S1B**). In addition, a principal component analysis (PCA) revealed clear clustering based on lipid profiles between early (16h) and late (48h) timepoints post viral infection (**Figure S1C).** PCAs also showed clustering with NIC treatment of virus infected cells at early (16h) and late timepoints (48h) (**Figures S1D and E, respectively**). Measurement of supernatant SARS-CoV-2 genome copy with RT-qPCR revealed a significantly higher quantity of released progeny genomes at 48h than 16h, indicating that more viral egress is occurring at the 48h timepoint, which is consistent with literature **(Figure S2)**. We did not perform any measurements of infectious progeny virions (i.e., plaque assay), so the antiviral effect of NIC in this experiment cannot be conclusively determined.

As SARS-CoV-2 infection caused perturbations in the global lipid profiles (**Figure 2A**), we next sought to characterize the modulations of different lipid classes between early (16h) vs late (48h) timepoints post viral infection. In total, we identified 720 lipids from 34 classes (**Table S2, S3**) in SARS-CoV-2 infected host cells covering all major lipid classes (**Figure 2A**). Phosphatidylcholine (PC), phosphatidylethanolamine (PE), plasmalogens (ether linked lipids) and triacylglycerol (TG) represented the most frequent lipid classes identified, which is expected given their abundance in cell membranes and lipid droplets (**Figure 2A, Figure 3**).

**Figure 1.**
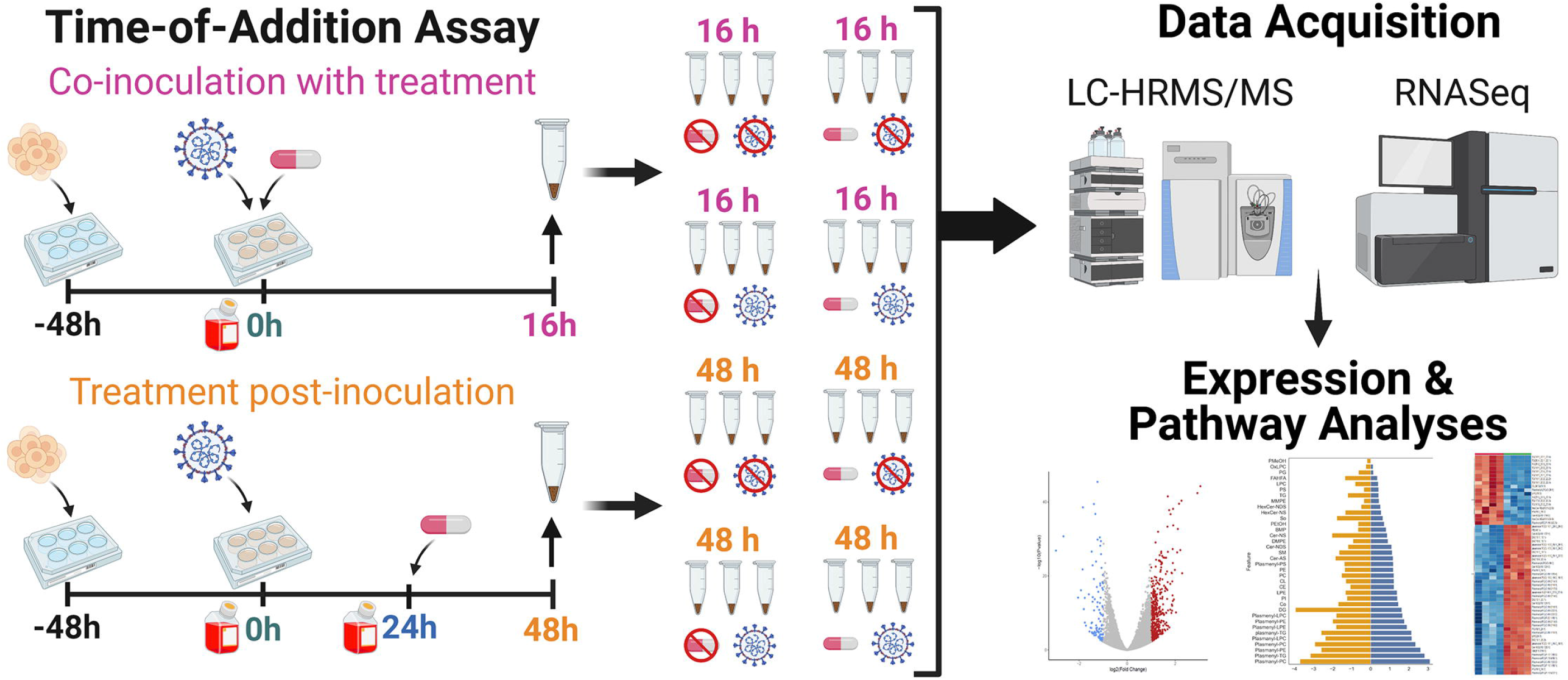
Workflow for lipidomic and transcriptomic profiling of Vero E6 cells after SARS-CoV-2 infection with and without niclosamide treatment. We used a time-of-addition assay experimental design to (i) capture the lipidomic profile of SARS-CoV-2 infected Vero E6 cells, and (ii) explore the effect of niclosamide on the lipidomic profile of Vero E6 cells when added in the absence of infection, with SARS-CoV-2 virus, or at 24h post-infection. Each replicate experimental condition (n=3) was processed for LC-HRMS/MS or RNASeq analyses. Samples were seeded 48h prior to the start of the experiment (t = 0h). For all samples, media was changed at the start of the experiment (t = 0h) and infected with virus (for infected sample groups). Sample collection is denoted by an up arrow and tube above the timeline, media changes are denoted by red bottles, addition of virus and DMSO/drug are denoted as well. Separate samples for each condition were collected for LC-HRMS/MS and RNAseq analysis, respectively.

**Figure 2.**
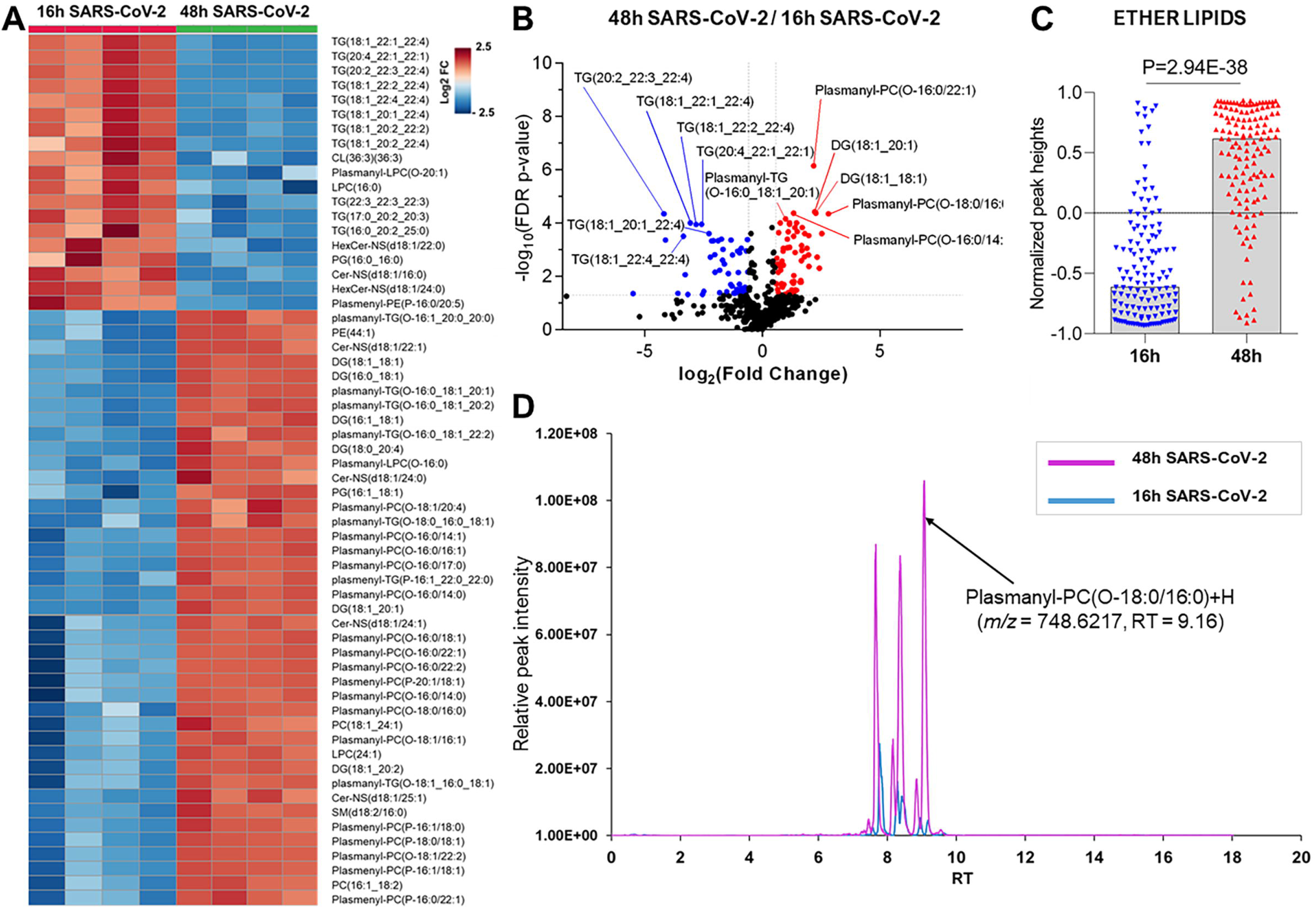
Global lipidomics analysis in SARS-CoV-2 infected Vero E6 cells at 16h and 48h. (A) Hierarchical cluster heatmap analysis depicting the major affected lipid clustering between 16h and 48h infection. (B) Volcano plot showing the differential lipid abundance with SARS-CoV-2 infection between 16h and 48h. The primary significant upregulated lipids were ether-linked (plasmanyl and plasmenyl). (C) Bar graph showing the total abundance of ether lipids between the time points. (D) Extracted ion chromatogram comparison showing relative peak intensity of a single ether lipid between 16h (blue) and 48h (magenta) infection. The arrow points to the lipid identified as Plasmanyl-PC (O-18:0/16:0)+H at retention time 9.16. Identification was confirmed by accurate mass and MS/MS.

**Figure 3.**
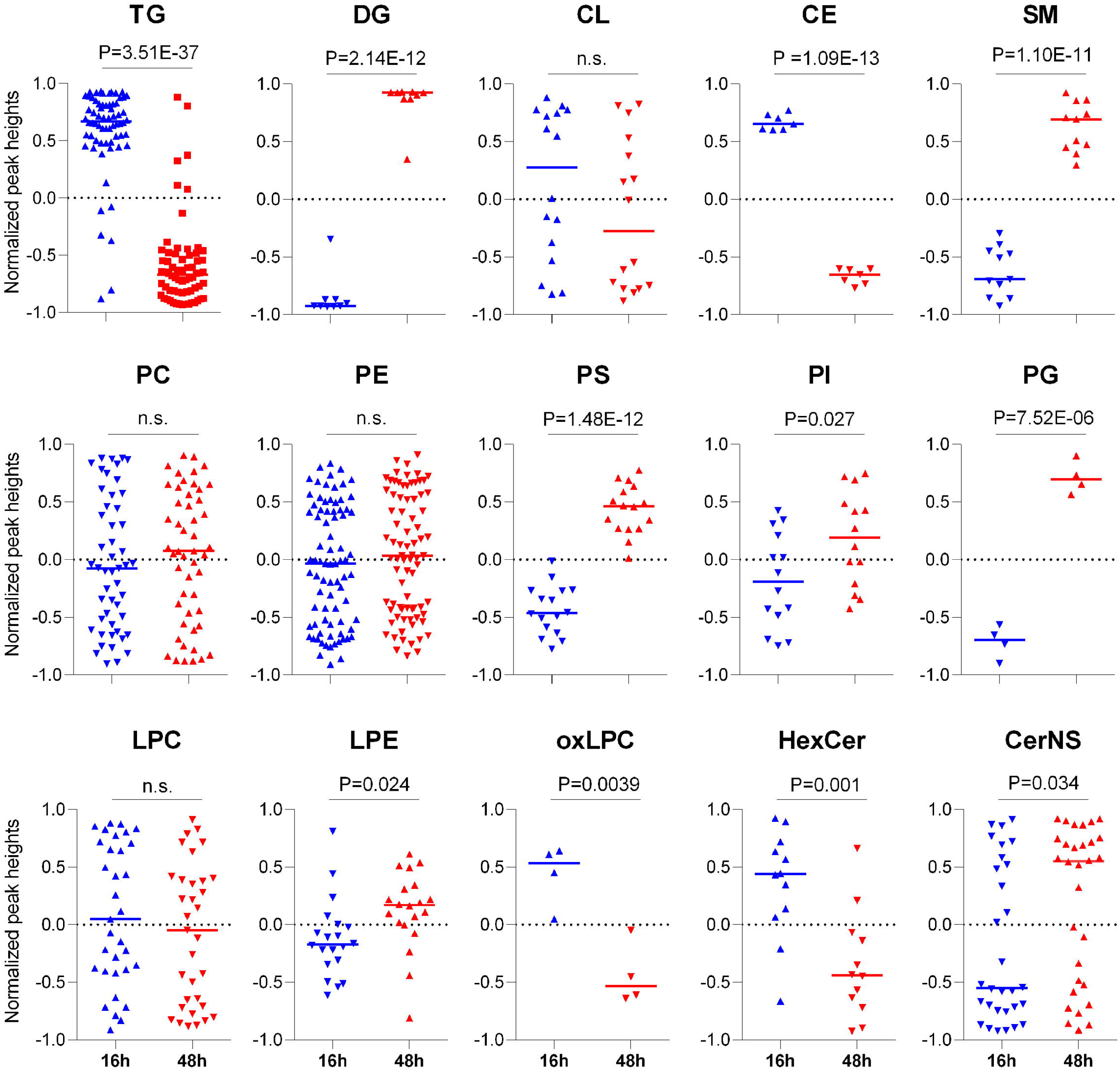
Comparison of individual lipid classes at early (16h) and late (48h) viral infection. FDR P-value based on T-test shown above each plot. Individual triangles represent each lipid detected and expression compared at both time points. Abbreviations: TG-triglycerides, DG-diglycerides, CL-cardiolipin, CE-cholesterol ester, SM-sphingomyelin, PC-phosphatidylcholine, PE-phosphatidylethanolamine, PS-phosphatidylserine, PI-phosphatidylinositol, PG-phosphatidylglycerol, LPC-lysophosphatidylcholine, LPE-lysophosphatidylethanolamine, oxLPC-oxidized lysophosphatidylcholine, HexCer-hexosylceramide, CerNS-ceramide with non-hydroxy fatty acid, n.s.-not significant.

Hierarchical clustering analysis of the differentially regulated lipids (FC>1.5, adjusted p-value <0.05) revealed specific lipid classes were associated with early or late events during productive virus infection (**Figure 2A, Table S2**). We found that ether lipids including plasmanyl-TG, plasmenyl-TG, plasmanyl-LPC, plasmanyl-PC, plasmenyl-PC, and plasmenyl-PE, as well as diacylglycerides were elevated at 48h post viral infection (**Figure 2B, Table S3**). We measured the relative abundance of the total ether linked lipids and observed a highly significant elevation (P-value = 2.94E-38) of total ether linked lipids at 48h post viral infection (**Figure 2C, Figure S3**), suggesting a potential role of ether linked lipids in SARS-CoV-2 pathogenesis. A comparative extracted ion chromatogram of plasmanyl-PC (O-18:0/16:0)+H, as an example of a specific ether lipid is shown (**Figure 2D**) to convey the clear difference in profiles between the two time points tested. On the other hand, TGs, cholesterol esters (CE), and hexosylceramides (HexCer) were elevated at early stages of infection (16h) (**Figure 2A and B, Figure 3, Figure S3, Table S3**). The repertoire of lipid classes that partitioned to early steps in virus infection (16h) and later stages of increased virus replication and egress (48h) are shown in **Figure 3**.

Lipids that contained longer chain fatty acids with higher degrees of unsaturation including TGs (4-9 double bonds (db)), PC (4-5 db), CL (5-6 db), PE (1-6 db), and PS (1-4 db) were elevated in early (16h) viral infection (**Figure 3, Table S3**), a trend consistent with what was described in SARS-CoV-2 infected golden Syrian hamsters (Rizvi et al., 2021). In contrast, saturated fatty acids (SFA), monounsaturated fatty acids (MUFA) and long chain fatty acids (LCFA) were found to be significantly downregulated in early (16h) viral infection but elevated at 48h post viral infection (**Table S3**), consistent with previous reports (Koyuncu et al., 2013; Nguyen et al., 2018). We identified changes in lipids that may be related to a cellular response to compensate for virus energy utilization, viral particle formation from lipid droplets, vesicle transport and autophagosome formation, noting a significant increase in plasmalogens (ether lipids), sphingolipids (SM and CerNS), glycerophospholipids (PI, PS and PG), lysophospholipids (LPE) and glycerolipids (DG) and a significant decrease in TG and CE lipids at 48h (**Figure 2A-B, Figure 3**).

We observed a differential regulation of lipids during increased levels of viral replication. Of note, we observed significant increases in PG, PS and PI, but not a significant change in PC or PE phospholipids. This differential regulation of phospholipids could suggest formation of membranes for the viral envelope as most mammalian cells are high in PC and PE phospholipids (Deng and Angelova, 2021). The significant decrease in TGs and CEs is likely related to virus utilization of lipid droplets to make viral particles needed for replication. Lipid droplets are used by cells to store neutral lipids that are utilized for energy needs, but viruses require these lipids for replication and thus hijack the lipid droplets to enable their own growth. The increase in plasmalogens is also associated with viral replication and increased levels have been detected in the serum of patients with ZIKV and other viruses (Cloherty et al., 2020; Mazzon and Mercer, 2014).

### Niclosamide treatment distinctly impacted ether lipids and this effect correlated with increased viral replication and virus egress

To understand these lipid modulations in the context of therapeutic potential, we used niclosamide (NIC), an anti-parasitic drug, which was found to suppress MERS-CoV propagation by enhancing autophagic flux through the Beclin1 (autophagy regulator/antiapoptotic protein)-SKP2 (S-phase kinase-associated protein 2) pathway (Gassen et al., 2019). NIC has also been proposed to be a potent candidate drug for repurposing in the treatment of SARS and COVID-19 (Gassen et al., 2021; Pindiprolu and Pindiprolu, 2020; Wu et al., 2004). The influence of SARS-CoV-2 infection on Vero E6 cell lipid metabolism, and inhibition of autophagic flux encouraged us to assess if NIC may impart therapeutic potential to SARS-CoV-2 infection.

We first examined the effect of NIC on Vero E6 cells in the absence of infection to establish a baseline for both the 16h and 48h time points (**Figure 4, Figure S4, Table S7)**. In total, we identified 520 lipids across all samples, with distinct profiles between DMSO control and NIC treated cells. We observed a profound reduction across both time points in the lipid profiles for TG, and this effect was more pronounced at the 48h time point (**Figure 4A**). In the absence of NIC treatment, plasmalogens (plasmanyl-PE, -TG, and -PC as well as plasmenyl-LPC, -PE, -LPE, -PC and -TG) were found to be differentially abundant at 48h as compared to 16h, which is expected given the cell growth during this time frame (**Figure 4A and Table S4**). However, at the 48h time point (24h post treatment) NIC reduced the abundance (>2 log2 fold-change) of HexCer-NDS, while in parallel, increased the abundance (>2 log2 fold-change) of plasmenyl- and plasmanyl-TG, PI, plasmanyl-PE and -PC, as well as BMPs (**Figure 4B and Figure S5).**

**Figure 4.**
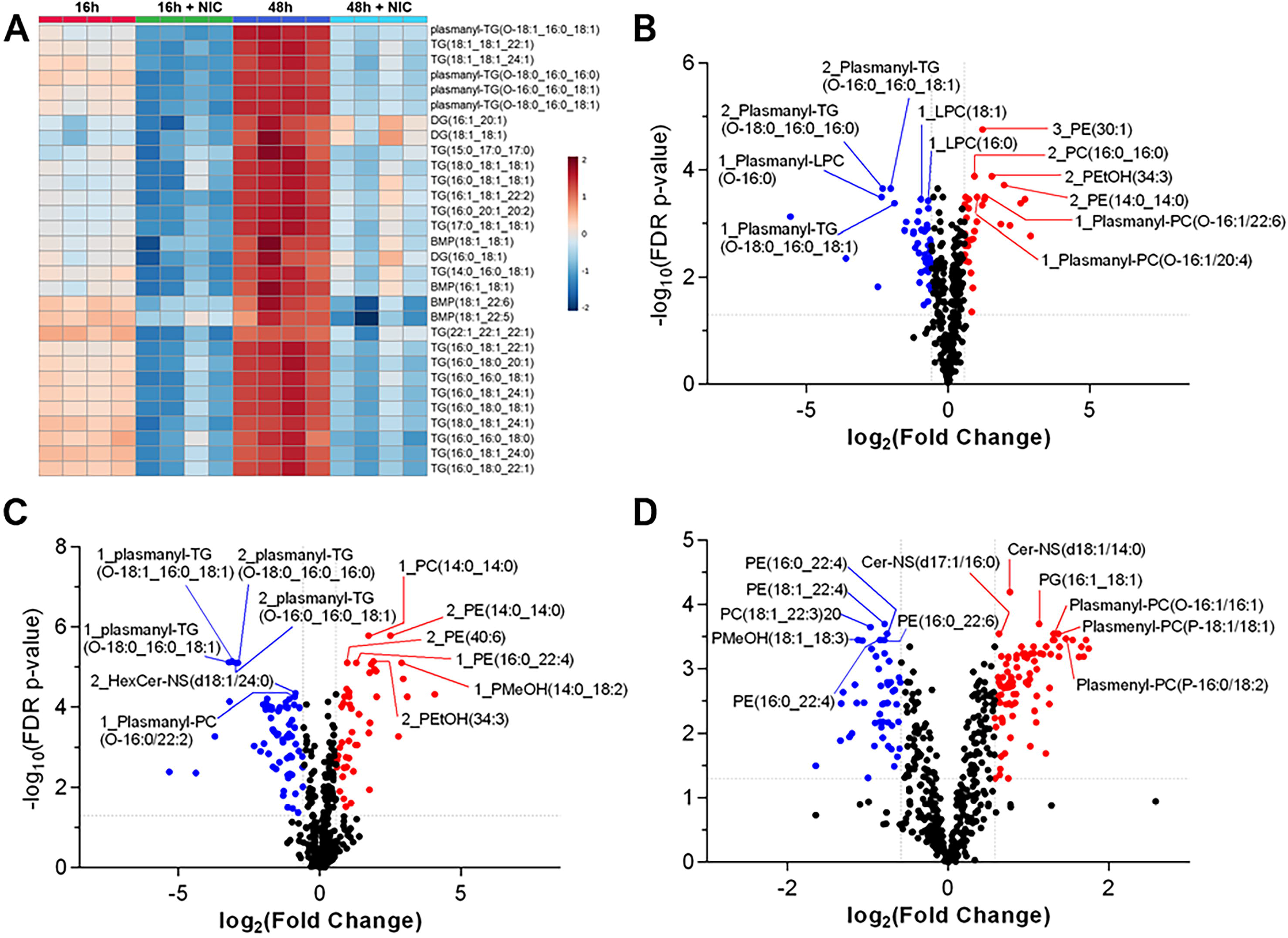
Niclosamide modulates lipid metabolism in Vero E6 cells in the absence of SARS-CoV-2 infection. Heatmap showing the changes in neutral lipids across the time points with and without NIC (A). Bar graphs of the significant lipids at 16h vs 48h (B) and 48h vs 48h + NIC (C). Fold change of less than 1.5 and p-value less than or equal to 0.05.

Having established a baseline effect of NIC on lipid metabolism in Vero E6 cells, we next sought to understand the effect of NIC on lipid metabolism during SARS-CoV-2 infection. We globally profiled lipids from cells infected with SARS-CoV-2 and treated with DMSO as compared to cells infected with SARS-CoV-2 and treated with NIC at 16h (**Figure S1B**) and 48h (**Figure 5A, Figure S1E**) using our UHPLC-HRMS approach. A PCA identified a clear separation between NIC treated and untreated SARS-CoV-2 infected samples (**Figure S1B, D and E**). We observed that the treatment of Vero E6 cells with NIC for 24h starting 24h after SARS-CoV-2 infection (48h) robustly impacted the global lipid profile as compared to NIC treatment beginning simultaneously with SARS-CoV-2 infection (**Figure 5B**), suggesting that the antiviral activity of NIC is also dependent on host cell state (in this case, infection as opposed to cell growth described above), which in turn correlated with higher levels of virus replication and virion production (**Figure S2**). However, no significant reduction in viral genome copy was observed in (NIC) treated cells compared to DMSO controls at either timepoint (**Figure S2**). This corroborates previous findings that NIC does not act to inhibit virus entry or genome replication (Gassen et al., 2021; Pindiprolu and Pindiprolu, 2020). NIC reduced DG as well as several plasmalogens, both of which would have otherwise been increased by SARS-CoV-2 infection at the 48h time point (**Figure 5A, Figure 2, Figure 3**). Strikingly, we observed that NIC treatment downregulated total ether lipids (P= 2.49E-38) (**Figure 5C**), suggesting that NIC may impart anti-viral effect via affecting ether lipid metabolism. As indicated earlier, we observed an increase in ether lipids at 48h in cells infected with virus alone, suggesting a key role of ether lipids in viral replication **(Figure 2C)**. Upon treatment with NIC, it appears that this lipid pathway is disrupted significantly, and thus may be affecting viral replication.

**Figure 5.**
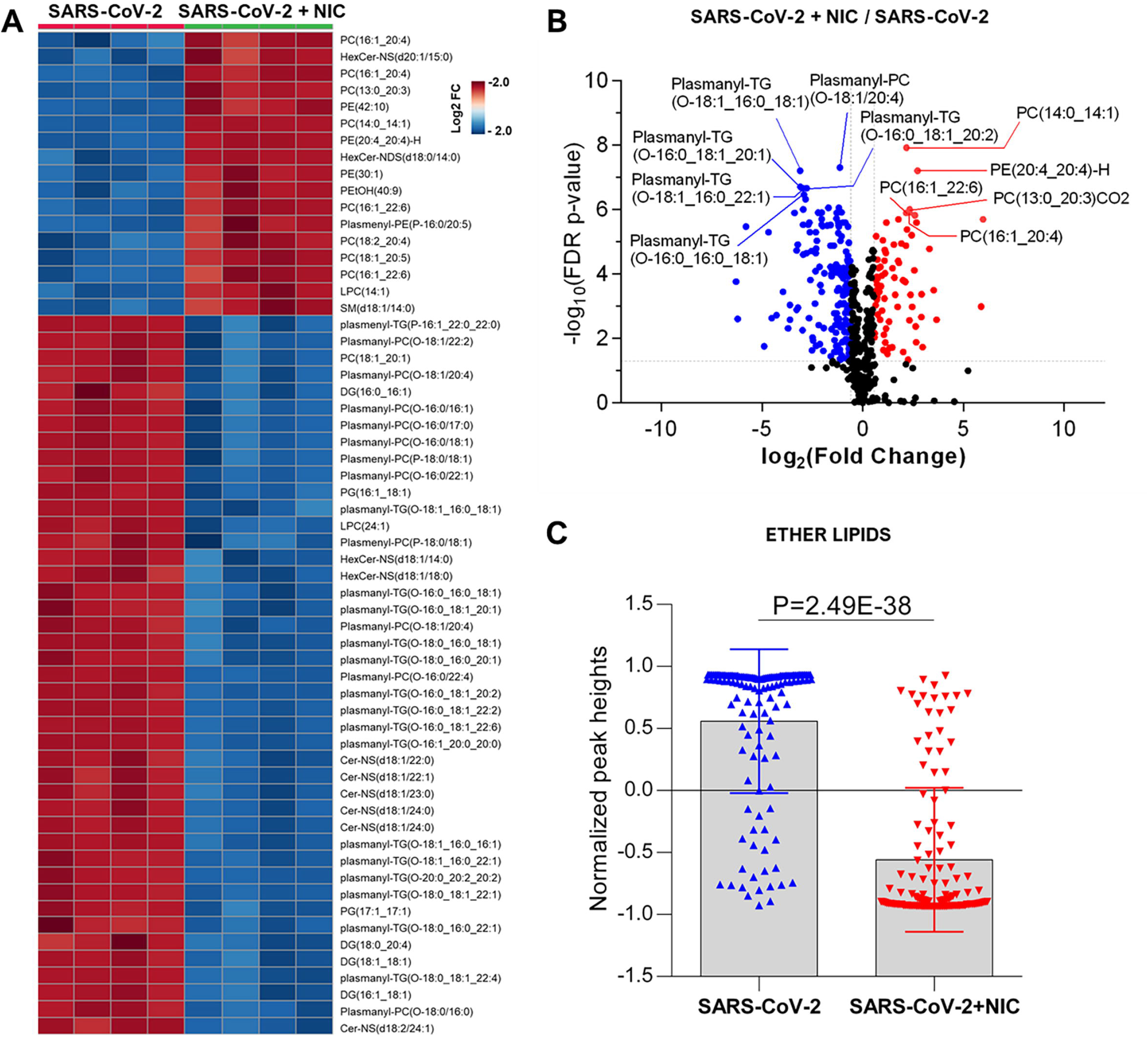
Niclosamide (NIC) impact ether-linked lipid production during viral replication at 48h. (A) Heatmap of the top 70 lipids (Fold change ≥ 1.5 and p-value ≤ 0.05). (B) Volcano plot showing the significant up and down regulated lipids with NIC at 48h. (C) The total ether lipids were down regulated with NIC treatment.

### Transcriptional profiling captures the broader cellular impact of virus infection and treatment with niclosamide

We assessed if virus-induced changes in lipidomic profiles corresponded to canonical transcriptional regulation of genes that are known to be involved in lipid metabolism, autophagy, phosphorylation, and vesicle transport. Furthermore, we explored if treatment with NIC alters this profile, and whether time of NIC addition alters the course of infection. To do this, we used RNASeq to capture the global cellular gene expression of i) Vero E6 cells over time (effect of time), ii) Vero E6 cells during cellular SARS-CoV-2 infection (effect of virus) and iii) following treatment with NIC (effect of drug).

We did not observe any significantly differentially expressed (DE) genes over time in our Vero E6 culture alone (0h vs 16h, 0h vs 48h, 16h vs 48h) (**Table S6**), suggesting while the cellular background of Vero E6 cells during these different time periods may be dynamic, the age of the cell culture did not confound other comparisons made in this study. Similarly, we did not observe any differences between cells treated with SARS-CoV-2 and NIC at 16h versus 48h. We did, however, observe differences with NIC treated cells at 16h versus 48h (860 DE genes, 474 up-regulated and 386 down-regulated at 48h) (**Table S1B, Table S6**), as well as SARS-CoV-2 infected cells at 16h versus 48h (109 DE genes, 48 up-regulated and 61 down-regulated at 48h) (**Table S1B, Table S6**).

SARS-CoV-2 infection resulted in significantly DE genes across all our comparison groups (**Table S1B, Table S5, Figure 6**. Viral infection resulted in 585 DE genes at 16h (474 up-regulated; 111 down-regulated), and 562 DE genes at 48h (433 up-regulated; 129 down-regulated) (**Table S5**). SARS-CoV-2 and NIC conditions resulted in more DE genes than SARS-CoV-2 infection alone; NIC and viral infection resulted in 3,083 DE genes (1,301 up-regulated; 1,782 down-regulated) at 16h, and 1,595 DE genes (706 up-regulated, 889 down-regulated) at 48h (**Table S5**). Overrepresented GO terms associated in both up- and down-regulated DE datasets contained a large number of lipid phosphate metabolism, lipid transport, autophagy and lysosome-associated terms. Terms related to phosphatidylinositol, phosphatidylethanolamine and phosphatidylglycerol were especially prevalent, mirroring observed changes in the lipidomics data. At 16h, viral infection induced an up-regulation of GABARAPL1, MAP1LC3A, PIK3C3, USP30-ORG3a and TBK1 (also up-regulated at 48h), and a down-regulation of STK11, LARP1, ZC3H12A, TFEB, TICAM1, GOLGA2, PIK3R2, IFI16, ULK1, WDR81, SQSTM1 (also down-regulated at 48h), and HK2. As per viral infection, NIC and virus treated cells (at 48h) also saw an up-regulation of GABARAPL1, MAP1LC3A, USP30-ORG3a, BMF, and TBK1; WDR81 was also down-regulated in the NIC and virus treatment at 48h.

**Figure 6.**
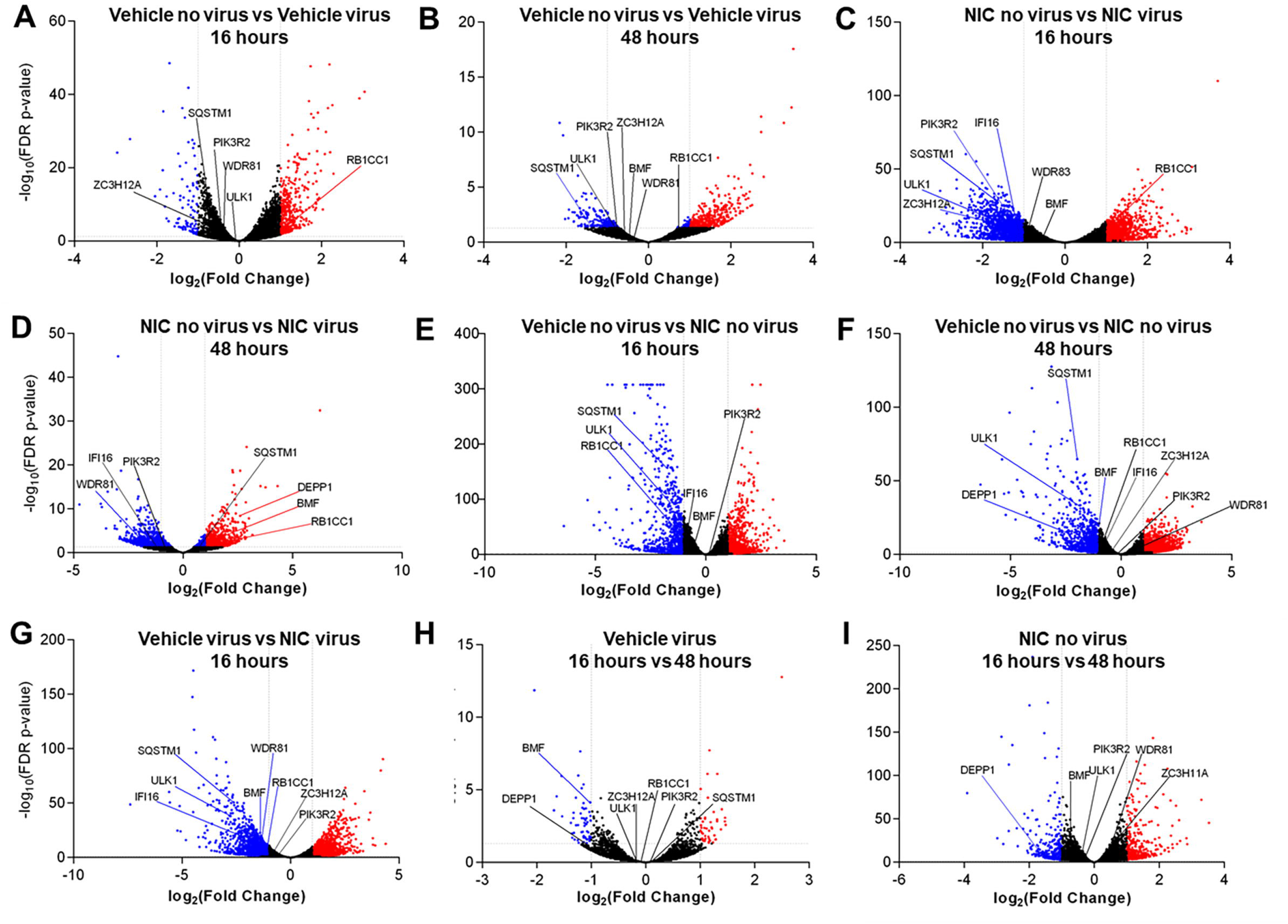
Significantly expressed genes across SARS-CoV-2 and Niclosamide conditions. Genes were designated as significantly expressed using DESeq2 when their Bonferroni adjusted P-value was ≤0.05, and the |log2(Fold Change)| was ≥1. Each dot represents an individual gene. Blue dots are significantly downregulated genes, red dots are significantly upregulated genes, and grey dots are not significantly differentially expressed genes. (A) DMSO treated no virus versus DMSO treated, virus samples at 16 hours, (B) DMSO treated no virus versus DMSO treated virus samples at 48 hours, (C) Niclosamide treated no virus versus niclosamide treated virus samples at 16 hours, (D) Niclosamide treated no virus versus niclosamide treated virus samples at 48 hours, (E) DMSO treated no virus versus niclosamide treated no virus samples at 16 hours, (F) DMSO treated no virus versus niclosamide treated no virus samples at 48 hours, (G) DMSO treated virus versus niclosamide treated virus samples at 16 hours, (H) DMSO treated virus at 16 hours versus 48 hours, (I) Niclosamide treated, no virus samples at 16 vs 48 hours.

There were significant transcriptomic differences noted at 16h and 48h with NIC treatment (**Figures 6E and F**), and at 48h with NIC treatment and SARS-CoV-2 infection (**Table S1B, Table S7, Figure 6G**. NIC treatment alone induced 2,447 DE genes at 16h (1,301 up-regulated; 1,146 down-regulated), and 2846 genes at 48h (1,707 up-regulated; 1,139 down-regulated), while viral infection and NIC treatment resulted in 3,617 DE genes at 16h (1,698 up-regulated; 1,919 down-regulated), although no DE genes were noted by 48h (**Figures 6E-G, Table S7**). Overrepresented GO terms with NIC treatment related to autophagosome assembly, regulation of autophagy, intracellular lipid metabolism, lipid localization and lipid homeostasis (**Figure S6**). Similar genes were noted for the effect of NIC as were with the effect of virus. RB1CC1 was up-regulated at 16h with virus infection (**Figure 6A**) but was down-regulated upon NIC treatment (**Figure 6E**) at 16h. Conversely, BMF was down-regulated with NIC at 48h, but was up-regulated upon NIC and virus treatment at 48 hours. Other genes that were down-regulated with viral infection (GOLGA2, IFI16, WDR81, HK2, SQSTM1, ULK1) remained down-regulated with NIC treatment (with and without virus).

## DISCUSSION

A virus subjugates host lipid molecules as a vehicle for entry, intracellular transport, virus replication, assembly of infectious viral particles, and egress (Miyanari et al., 2007). Ether lipids have been identified as an important lipid class for efficient membrane trafficking, endocytosis, transcytosis, and internalization of particles (Thai et al., 2001; Deng and Angelova, 2021). Deficiency of ether lipids in several model systems affects plasma membrane function, as well as structural changes in the ER and Golgi cisternae (Bazill and Dexter, 1990; Gorgas et al., 2006). SARS-CoV-2 infection has demonstrated a diverse range of COVID-19 disease severities that correlate with increased viral replication (Fajnzylber et al., 2020). Unlike during the early infection stage at 16h, where virus load and replication is low, we observed that SARS-CoV-2 infection uniquely elevated ether linked lipids after 48h of infection, a phase that represents increased virus replication, virus assembly and egress as well as host cell energy utilization (Abu-Farha et al., 2020; Dimitrov, 2004; Mazzon and Mercer, 2014). Increased viral replication drives infection severity and may lead to a perturbation of the host cellular molecular landscape (Fajnzylber et al., 2020).

We observed that NIC treatment significantly downregulates the ether lipid profile during infection. We also observed that NIC treatment in Vero E6 cells causes a reduction in TG content at both 16h and 48h suggesting a disruption of lipid droplet formation. (**Figure 4A**). NIC has previously been tested as an anti-obesity drug, and a study using it associated with a high fat diet showed a decrease in total TGs (Al-Gareeb et al., 2017). NIC also affects several lipid species, including PE, which may stimulate the autophagy process and inhibit ether lipid elevation during viral propagation. However, NIC has dual activity in cells and is capable of influencing autophagic processes through canonical and non-canonical pathways (Liu, et al., 2019). Whereas in uninfected cells, NIC reduces autophagic flux as well as the global lipid profile (causing a decrease in plasmalogens, TGs and PE lipids), in infected cells, NIC induces autophagy and the subsequent reduction of the ether lipid profile. NIC has been shown to affect the function of lysosomes by preventing the acidification of the vacuole (Jung et al., 2019). This disruption of lysosomal function results in increased autophagic flux similar to the likely primary mechanism of action underlying the broad antiviral activity of chloroquine (Mauthe et al., 2018). Importantly, we identified that ether linked lipids were only affected by NIC treatment relative to productive virus infection at 48h, suggesting that the activity of NIC may be dependent on the cellular state or level of virus burden and virus replication kinetics.

To understand the critical lipids related to autophagy activation, we analyzed the phosphatidylethanolamine (PE) levels in NIC treated SARS-CoV-2 infected cells. PE is one of the central lipids that conjugates the factors for induction of autophagy through the formation of the autophagosome (Xie et al., 2020). We observed that the relative abundance of PE was significantly elevated with NIC treatment at 48h while there was no impact of NIC on PE levels at 16h, suggesting that NIC enhanced the autophagy machinery upon viral replication to directly affect downstream virus assembly, trafficking, and egress. Different signaling lipids such as PS, PI, and PGs play critical roles in endosome trafficking and maturation (Mazzon and Mercer, 2014) and help in infectious viral particle transport. Our study first identified that NIC treatment significantly reduced the global level of different glycerophospholipids such as PS, PI, and PG at 48h, but had no impact on early infection at 16h.

TGs are the major form of lipids enriched in lipid droplets. Drugs that stimulate TG downregulation usually activate autophagy and may prevent a cell from viral infection and pathogenesis. We identified that NIC treatment significantly reduced global levels of TGs in SARS-CoV-2 at 48h treatment, suggesting that NIC treatment potentiates TG breakdown in lipid droplets to help reduce viral replication. We also found that NIC treatment showed either no effect or very low impact on lipids that were involved in lipid droplet formation at 16h. In fasting conditions, lipid droplets are utilized for fatty acid oxidation to produce energy as intact TGs are broken down to release fatty acids in a process called lipophagy. The breakdown of TGs via hydrolysis is mediated by autophagy and we observed a decrease in TGs with NIC treatment. Overall, these findings demonstrate that NIC treatment likely activates autophagy and consequently impacts ether linked lipids and other lipids needed for viral infection and propagation.

Bis(monoacylglycerol)phosphates (BMP), identified as a new lipid of endosome-derived extracellular vehicles (EVs), also act as cholesterol transporters in cooperation with other factors and facilitate viral infection and replication (Luquain-Costaz et al., 2020). Interestingly, our study detected three different forms of BMP and two different forms of cholesterol esters (CE) that were all significantly downregulated with NIC treatment. Besides the impact of NIC on different autophagy related lipids, we also investigated the role of NIC on bioactive lipid metabolism and apoptosis. We observed a dichotomous pattern of CerNS and SM lipids, where CerNS decreased after NIC treatment (48hV vs 48hV+NIC) while SM were increased, indicative of cell death or cell survival, which may be partly explained by the cell state-dependency of NIC that we have observed in the presence or absence of SARS-CoV-2 infection in Vero E6 cells. Interestingly, we identified a notable reduction of gangliosides detected as HexCer (which are also called glycosphingolipids) upon NIC treatment at 48h in both uninfected and SARS-CoV-2 infected Vero E6. Gangliosides have been implicated in the induction of autophagic death in mammalian cells (Hwang et al., 2010), suggesting that NIC can increase cell survival at homeostasis as well as under stress/infection conditions. This activity may in part contribute to the low cytotoxicity for this drug for several different cell lines, with the reported CC_50_= 50 µM (Jeon et al., 2020; Xu et al., 2020). Although gangliosides have also been shown to bind cooperatively with ACE-2 to the SARS-CoV-2 spike protein (Fantini et al., 2021), our experimental design did not include a pretreatment of cells prior to virus infection. The search for highly safe drugs for prophylaxis against COVID-19 remains elusive and controversial, but these data may suggest a role for niclosamide for future outbreaks given its broad antiviral activities. Considering the potential link between plasmalogens and the observed cytokine and lipid storms (Deng and Angelova, 2021) in severe COVID-19 patients, NIC may offer to a two-pronged treatment approach, i.e., reduce virus abundance and provide a check on an uncontrolled inflammatory response induced by SARS-CoV-2.

These findings are further supported by the global transcriptome profile that we captured for each of the treatments/conditions, wherein hallmarks of and the induction of autophagy related genes (such as DEPP1 (an autophagy regulator) (Salcher et al., 2017)), several lipases (Gassen et al., 2021), as well as genes implicated in the induction of apoptosis and genes that are markers of cell proliferation. Furthermore, GO overrepresentation analyses showed significantly upregulated terms such as death receptor activity, MAP kinase phosphatase activity, tumor necrosis factor-activated receptor activity, and phosphatidylinositol kinase activity. To investigate the role NIC has on the reversal of SARS-CoV-2 infection induced autophagy dysregulation, we explored the function of several genes that were differentially expressed. RB1CC1 (RB1 inducible Coiled-Coil 1) along with ULK1 are part of the ULK complex that initiates and regulates autophagy (Zachari and Ganley, 2017). RB1CC1 has also been shown to influence viral infection (Mauthe and Reggiori, 2016), as a depletion of RB1CC1 led to an increase of encephalomyocarditis virus replication. Here, RB1CC1 was up-regulated by SARS-CoV-2 at 16h; however, was significantly down-regulated with treatment of NIC at 16h. We also observed down-regulation of ULK1 in NIC treated cells at 16h. This elucidates the potential role of RB1CC1 and ULK1 in early infection with SARS-CoV-2 and illuminates how NIC may regulate autophagy. Interestingly, BMF was down-regulated at 48h with NIC, but up-regulated upon NIC treatment of virus infected cells at 48h. BMF encodes for a Bcl-2-modifying factor that is responsible for apoptotic regulation and has been reported as a target facilitating viral evasion (Zamaraev et al., 2020). Additionally, autophagy genes IFI16 (Duan et al., 2011; Jiang et al., 2021; Kim et al., 2020; Wichit et al., 2019), ZC3H12A (Srivastava et al., 2020), SQSTM1 (Gassen et al., 2021), WDR81 (Zhu et al., 2021), and PIK3R2 (Weisberg et al., 2020) are associated with viral infection, including SARS-CoV-1 (Gassen et al., 2021; Srivastava et al., 2020) and were downregulated in this study when cells were treated with NIC. Taken together, this provides evidence that SARS-CoV-2 infection leads to dysregulation of autophagy, and that NIC acts to reverse this effect.

Several omics studies have made a significant contribution to the identification and understanding of potential antiviral therapies for COVID-19 (Mahmud and Garrett, 2020). Bioactive molecules including PUFA lipids such as oleoylethanolamide (OEA), arachidonic acid (AA), eicosapentaenoic acid (EPA), and docosahexaenoic acid (DHA) are known from existing viral pathogenesis to inactivate enveloped viruses and inhibit pathogen proliferation/replication (Das, 2020; Ghaffari et al., 2020; Tam et al., 2013). Accumulation of lipids including sphingolipids such as ceramides has been shown to negatively affect viral pathogenesis (Soudani et al., 2019; Young et al., 2013). Lipid metabolism; specifically catabolism, biosynthesis, and peroxidation play critical roles in autophagy or apoptosis-mediated cellular homeostasis including cell survival and death (Xie et al., 2020). Autophagy can act as an anti-viral or pro-viral mechanism; however, most viruses are found to inhibit autophagy signaling (Jackson, 2015). In the case of SARS-CoV-2, we know little about the relationship of virus infection with autophagy signaling. Importantly, we identified elevated levels of ether lipids (plasmanyl and plasmenyl) with increased viral replication, which was reported as a key lipid class for efficient membrane trafficking (Thai et al., 2001). We revealed that lipids are critical molecular factors for SAR-CoV-2 infection, entry, and viral replication into host cells. These lipids, which were elevated by SARS-CoV-2 infection may be counteracted by PUFA and VLCFA that the virus suppresses upon infection and entry (Kang et al., 2019).

Collectively, these data provide new insight into the mechanism of action for the antiviral activity of NIC, namely induction of lipophagy through the induction of autophagic pathways (**Figure 7**), which expands as well as refines our understanding of the pleiotropic antiviral mechanisms described for NIC (Braga, et al., 2021; Gassen, et al., 2021; Kao, et al., 2018). We can mechanistically infer that SARS-CoV-2 infection wrests control of cellular autophagy to ensure a productive infection of the host cell. Therapeutically, we demonstrated that NIC treatment stimulated autophagy by elevating PE levels and in parallel, downregulated total ether lipids, TGs, and other lipid molecules. Reduction of these key lipids by NIC treatment may lead to inhibition of SARS-CoV-2-induced signaling, which in turn drives viral endocytosis, vesicular trafficking, propagation, and viral egress. Activating autophagy using small molecule inhibitors/activators, oxysterols, or peptides is a promising approach for treating viral infections (Rubinsztein et al., 2012). However, most of the potential treatments (including NIC) suffer translational obstacles due to adverse side effects, inefficient drug delivery to target tissues, and poor bioavailability (Skotland et al., 2015). Although NIC is considered an essential medicine and is well tolerated, the long history of dosing safety is based on *per os* delivery and anti-parasitic activity within the gastrointestinal tract. SARS-CoV-2 infection begins in the lungs but has been shown in several clinical studies to have a cosmopolitan infectivity profile, including the brain, heart, and gastrointestinal tract. Notably, several repurposed drugs for COVID-19 were found to be associated with autophagy induction (Kim et al., 2012), raising the possibility that some drugs that are already in clinical use for the treatment of parasitic or bacterial infections may be acting, at least in part, via autophagy. The recent clinical testing of intranasal or inhaled NIC (Backer et al., 2021) suggests that the potential of this drug for use in mitigating COVID-19 severity is profound, compeling expansive exploration of different approaches to increase targeted delivery and efficacy.

**Figure 7.**
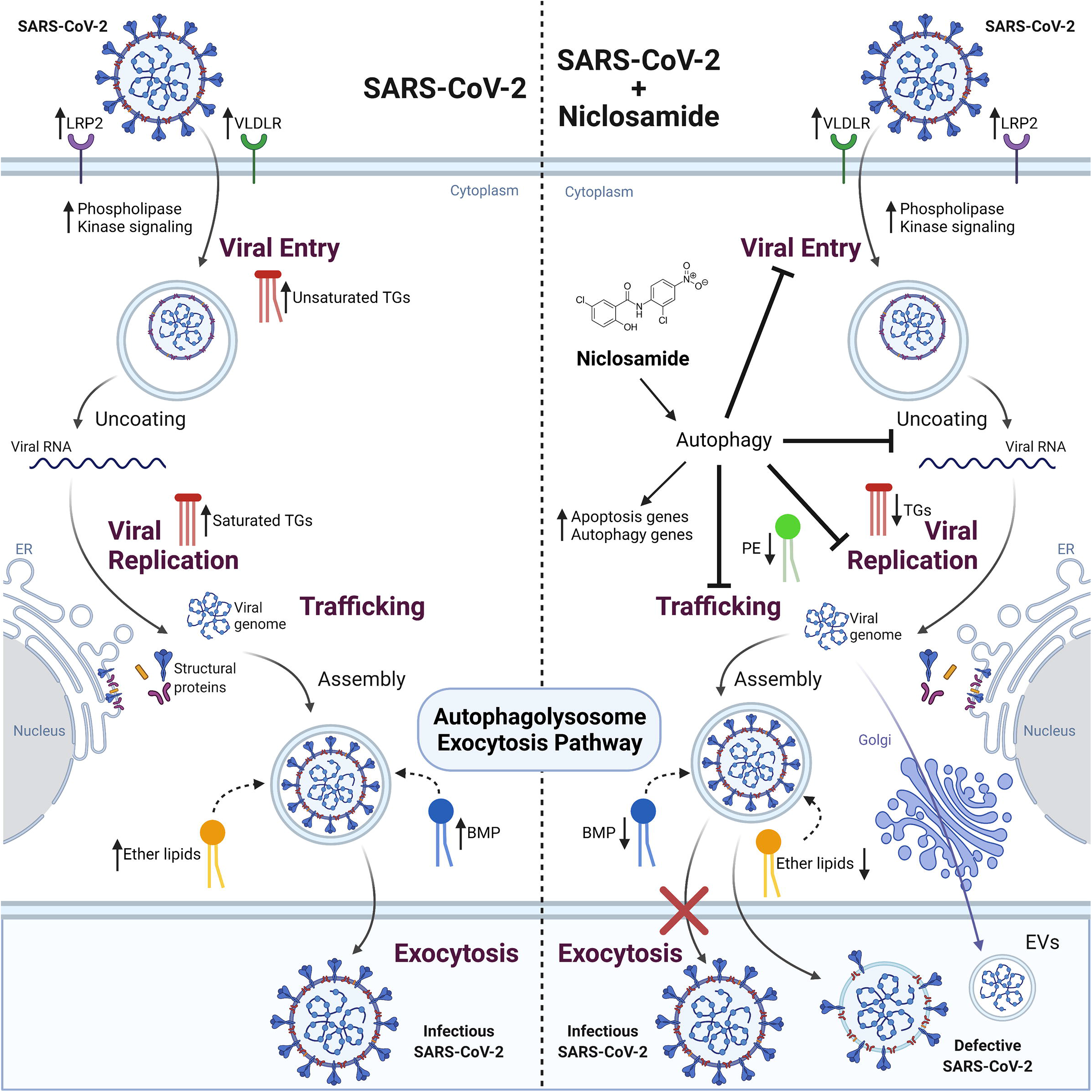
The effect of SARS-CoV-2 infection and NIC treatment on host cell lipid metabolism. SARS-CoV-2 infection in Vero E6 cells alters host cell lipid metabolism during early and late stages of infection. Increased transcription of lipid receptors LRP2 and VLDLR, and phosphorylation signaling regulators are observed throughout viral infection. Changes in TG composition from unsaturated to saturated acyl-chains occurs as a function of viral replication, with an overall decrease in TG lipids at late infection timepoints. This change corresponds with an increase to DG and BMP lipids that is indicative of energy consumption and incorporation into membranes and vesicles, activation of autophagy pathways, as well as impacting viral replication. Treatment of cells with NIC alters lipid composition and gene regulation corresponding to apoptosis and autophagy related pathways. Decreases to ether lipids (TGs and DGs) and BMP are observed and reflect a decrease to exocytosis pathways for viral egress.

### Limitations

Although we used a *C. sabaeus* kidney epithelial cell line (Vero E6) and a SARS-CoV-2 strain isolated from a Floridian patient, we observed a similar (in direction and magnitude) transcriptomic response to other SARS-CoV-2 transcriptomic studies, despite differences in MOI and experimental sampling timepoints. These changes were consistent across another Vero E6 study noting cell stress and apoptosis (DeDiego et al., 2011), as well as other cell systems, including human lung cells (Wyler et al., 2021), cardiomyocytes (Sharma et al., 2020), adenocarcinomic human alveolar basal epithelial cells (Daamen et al., 2021), and bronchial epithelial cells (Yoshikawa et al., 2010), infected with different betacoronaviruses, suggesting the responses detected herein may represent a core set of host SARS-CoV viral response genes, and that these genes are not exclusive to our study system. Our work is limited to an *in vitro* model; however, it was recently shown in a multiomics study of COVID-19 patient samples that the lipidomic profile can effectively partition COVID-19 disease severity (Overmyer et al., 2021); implying that the biology captured in our culture model is relevant *in vivo*. While these observations highlight potential utility of niclosamide, these data alone do not support its indication in the clinic in the treatment of COVID-19.

## STAR METHODS

### RESOURCE AVAILABILITY

#### Lead Contact

Further information and requests for resources and reagents should be directed to and will be fulfilled by the lead contact, Dr. Timothy Garrett.

#### Materials availability

This study did not generate new unique reagents.

#### Data and code availability

The raw and processed data generated here have been deposited in publicly accessible databases; the RNA sequencing data is available through NCBI’s GEO repository (Accession: GSE178157), and the lipidomics data is accessible via MetaboLights (Accession: MTBLS2943).

This paper does not report original code.

Any additional information required to reanalyze the data reported in this paper is available from the lead contact upon request.

### EXPERIMENTAL MODEL AND SUBJECT DETAILS

#### Vero E6 cells

Vero clone E6 cells (ATCC: CRL-1586) were obtained from Dr. Pei-Yong Shi (University of Texas Medical Branch). Vero E6 cells are an immortalized kidney epithelial cell line obtained from the African green monkey (*Cercopithecus sabeus*). The cells are female.

Cells were maintained at 37 °C and 5% CO_2,_ in Dulbecco’s Modified Eagle Medium (ThermoFisher #11965118) supplemented with 10% heat-inactivated fetal bovine serum (FBS), 1X L-glutamine (Gibco #25030-081), and 1X Penicillin/Streptomycin (Corning #30-001-CI). These cells have been authenticated.

#### SARS-CoV-2

SARS-CoV-2/human/USA/UF-13/2020 (GenBank: MT620766.1) was passaged in Vero E6 cells, with culture conditions as described above except using reduced serum media with 3% FBS rather than 10% FBS. Virus was collected after 4 days to establish a low passage virus stock within our biosafety level 3 (BSL-3) laboratory at the Emerging Pathogens Institute.

### METHOD DETAILS

#### Cell culture, infection, and niclosamide treatment

Cells were seeded in 6 well plates at a density of 500,000 cells/well. Each replicate sample represents an individual plate well, with three biological replicates per condition, per time point. The investigators were not blinded to the conditions. At hour zero (48 hours post seeding), media was removed and replaced with reduced serum Dulbecco’s Modified Eagle Medium (supplemented with 3% FBS, 1X L-glutamine, and 1X Penicillin/Streptomycin) to facilitate infection.

One vial of SARS-CoV-2 stock was thawed and diluted to a titer of 2×10^7^ plaque forming units (PFU) per mL. For infected conditions, virus was inoculated into each well with 12.5 µL of diluted virus stock for a multiplicity of infection (MOI) of 0.5 infectious virus particles/ Vero E6 cell. Niclosamide (MedChemExpress #HY-B0497, [NIC]) stock was dissolved in molecular grade DMSO and added to the indicated wells for a final concentration of 5 µM (0.5% v/v DMSO final concentration). An equivalent amount of molecular grade DMSO was used as a vehicle control. This concentration of NIC was selected to have antiviral effect but be well below the observed 50% cytotoxic concentration (CC_50_) of the drug (50µM) to avoid inadvertent cell lysis in treated conditions (Wu et al., 2004).

Samples were prepared and collected following a time of addition experimental design (Daelemans et al., 2011) as follows (**Figure 1, Table S1A-B**). The “0 hour” (0h) samples were collected 48h after cell seeding. The “16 hour” (16h) conditions were infected with virus, treated with NIC or DMSO at hour zero, and harvested at hour 16 of the experiment (**Figure 1**). The “48 hour” (48h) conditions were infected with virus at hour zero, supernatant was removed after 24h, and cells were treated with NIC or DMSO in fresh media. Cells were harvested at hour 48. Uninfected samples (with and without drug treatment) were also prepared and collected as above. For samples from infected conditions, the supernatant was removed at the indicated harvest timepoint, taking care not to disturb the cell monolayer, and retained at −80°C. The cells were washed once with 1X PBS. The PBS wash was removed and 0.25% trypsin-EDTA (ThermoFisher #25200056) was added to each well. The plates were then incubated at 37°C and 5% CO_2_ for 5-10 minutes until cells had completely detached from the cell surface. Trypsinization was halted by the addition of 1 mL FBS. The supernatant was removed, the cell pellet was resuspended in 1X PBS, and the sample was centrifuged at 700 RPM for 5 minutes. Supernatant was aspirated, and 100% methanol was added to each sample purposed for lipid analysis. Alternatively, 750 µL of TRIzol reagent (ThermoFisher #15596026) was added to each sample purposed for RNASeq. All samples were stored at −80°C until analysis. For uninfected conditions, harvest proceeded the same as described above except trypsinization was halted using 4 mL of complete Dulbecco’s Modified Eagle Medium instead of FBS, and pellets were flash frozen in liquid nitrogen instead of being resuspended in a chemical inactivation agent after the last centrifugation step. All samples were stored at −80°C until analysis.

#### Global lipid extraction

Cell pellets from each time point and condition were extracted using a modified Folch biphasic extraction procedure (Ulmer, et al. 2017). Samples were pre-normalized to the protein concentration (800 µg/mL) obtained using a Qubit 4.0 Fluorometer (Thermo Fisher Scientific). Lipids were extracted using ice cold 4:2:1 chloroform:methanol:water (v:v:v) containing 20 μ L of a 10X diluted internal standard mixture (stock solution of 50 ppm, w: v), and the organic phase was collected after centrifugation at 3260 x*g* for 5 min at 4°C, dried under nitrogen gas, and reconstituted in 75 μL of injection standard mixture (100 ppm, w:v). Internal standards used in this analysis covered a range of lipid classes and structures including: lysophosphatidylcholine (LPC 17:0), phosphatidylcholine (PC 17:0/17:0), phosphatidylethanolamine (PE 15:0/15:0), phosphatidylserine (PS 14:0/14:0), phosphatidylglycerol (PG 14:0/14:0), ceramide (Cer d18:1/17:0), diacylglycerol (DG 14:0/14:0), and sphingomyelin (SM d18:1/7:0). For injection standards, triacylglycerol (TG 17:0/17:0/17:0), LPC 19:0, PE 17:0/17:0, PS 17:0/17:0, and PG 17:0/17:0 were used. Except for TG, all other lipid standards were purchased from Avanti Polar Lipids (Alabaster, AL) while TG was purchased from Sigma Aldrich. All lipid standards were diluted prior to analysis in 1:2 (v/v) chloroform:methanol (CHCl_3_:MeOH) and a working standard mix was then prepared by diluting the stock solution with the same solvent mixture.

#### LC-MS-based lipid data collection and analysis

Ultra-high-pressure liquid chromatography coupled to high resolution mass spectrometry (UHPLC-HRMS) was used for data collection. Chromatographic separation was achieved using reversed phase chromatography (Dionex Ultimate 3000 RS UHLPC system, Thermo Scientific) with a Waters Acquity C18 BEH column (2.1 × 50 mm, 1.7 μm) (Waters, Milford, MA, US). The mobile phases consisted of solvent A (60:40 acetonitrile:water) and solvent B (90:8:2) isopropanol:acetonitrile:water), both containing 10 mM ammonium formate and 0.1% formic acid. The flow rate was 500 μL/min and the column temperature was maintained at 50°C. A multi-step gradient was used for separation starting with 0% B from 0-1 min, increasing to 30% B from 1-3 min, then up to 45% B from 3-4 min, 60% B from 4-6 min, 65% B from 6-8 min, held at 65% B from 8-10 min, increased to 90% B from 10-15 min, then increased to 98% B from 15-17 min and finally held at 98% B from 17-18 min before returning to initial conditions to equilibrate for the next injection. Samples were analyzed in positive and negative electrospray ionization as separate injections on a ThermoScientific Q-Exactive high resolution mass spectrometer (Thermo Scientific, San Jose, CA). Lipidomics data were compiled and annotated using LipidMatch (Koelmel et al., 2017).

#### RNA extraction

For total RNA extraction from flash frozen cells, the cell pellet was thawed on ice. Cell pellets were resuspended in 1 mL Trizol, and 200 µL chloroform was added to each sample. Samples were mixed thoroughly by vortexing, then incubated at room temperature for 3 minutes. Samples were centrifuged at 12,000 x*g* for 15 minutes at 4°C. The upper aqueous layer was transferred to a new tube and 500 µL isopropanol was added and mixed by vortexing, prior to incubation at room temperature for 10 minutes. RNA was pelleted by centrifuging at 12,000 x*g* for 10 minutes at 4°C, and supernatant was discarded. The pellet was rinsed in 75% ethanol and pelleted by centrifuging at 7,500 x*g* for 5 minutes at 4°C. The ethanol wash and spin were repeated a second time. The pellet was air dried in an inverted tube. Genomic DNA was digested using the TURBO DNA-free™ Kit (Invitrogen #AM1907) according to manufacturer instructions. Samples were mixed with 350 µL buffer RLT from the RNeasy® Mini kit (Qiagen #74104) and 250 µL 100% ethanol, then transferred to a RNeasy® Mini kit spin column. Columns were centrifuged at 8,000 x*g* for 15 seconds at room temperature, and flow-through was discarded. Buffer RPE (500 µL) was added to the column, which was centrifuged at the same conditions, and flow-through was discarded. The RPE wash was repeated a second time. The column was transferred to a fresh collection tube and centrifuged at 12,000 x*g* for 1 min to dry. RNA was eluted by adding 50 µL of nuclease free water and centrifuging for 1 minute at 8,000 x*g* and was immediately frozen at −80°C. To extract viral RNA from culture supernatant, supernatant was thawed, 200 µL of supernatant was mixed with 200 µL DNA/RNA Shield, and RNA was immediately extracted using the Quick-DNA/RNA™ Viral MagBead kit (Zymo Research #R2140), according to manufacturer instructions and then immediately frozen at −80°C.

#### Real-time RT-qPCR

To quantify viral genome copy in supernatant, one step real time reverse transcription qPCR (RT-qPCR) was performed using 4x Quantabio UltraPlex 1-Step ToughMix No Rox (VWR #10804-946) and CDC 2019-nCoV_N1 (nucleocapsid) primer and probe mix (see Key Resources Table for primer and probe sequences) at a final concentration of 22.5 µM, in a 20 µL reaction volume with 5 µL template. All samples were run in technical duplicates. RT-qPCR was run on a BioRad CFX96™ Real-Time System. The thermal cycling conditions were as follows: 50°C for 20 minutes, 94°C for 2 min; followed by 45 cycles of 94°C for 15 seconds, 55°C for 30 seconds, and 68°C for 10 seconds. The samples were quantified using a standard curve generated using a 2019-nCoV_N_Positive Control plasmid (IDT # 10006625). Standard curve points were plotted in Microsoft Excel v.2102 and cycle threshold values of unknown samples were determined from the logarithmic line of best fit equation to calculate genome copies/mL.

#### RNASeq analysis

All 27 RNA samples (see **Table S1A-B**) were directly sent to Novogene (https://en.novogene.com/), where in-house RNA quality control and library preparation was performed. Libraries were sequenced on an Illumina NovaSeq6000 platform using a 150bp kit with paired-end read mode.

#### Mapping, expression, and pathway analyses

Bioinformatic processing was completed using Galaxy (https://galaxyproject.org, (Batut et al., 2018)). Quality control was performed with Cutadapt v 1.16 (Martin, 2011) and FastQC v 0.11.8 (http://www.bioinformatics.babraham.ac.uk/projects/fastqc/) to remove adapters, <20 nucleotide reads and low-quality reads. RNA STAR v2.7.7a (Dobin et al., 2013) was used to map samples to the *Chlorocebus sabaeus* genome and associated annotation (GenBank accession # GCA_015252025.1). featureCounts v 2.0.1 (Liao et al., 2014) was used with Infer Experiment v2.6.4.1 (Wang et al., 2012) to determine the strandedness of the samples, and sum reads for each gene. Read counts for each sample were input into DESeq2 v 1.22.1 (Love et al., 2014) to call differential gene expression, analyzing the effect of time, drug, and virus on the samples. Resultant differential expression files were manually annotated using a combination of NCBI (O’Leary et al., 2016) and Ensembl (Yates et al., 2020) due to the poor annotation quality of the genome (https://www.ncbi.nlm.nih.gov/genome/annotation_euk/Chlorocebus_sabaeus/100/). Significantly regulated gene names were separately parsed out for each comparison and uploaded to ENRICHR (Kuleshov et al., 2016) for downstream gene ontology (GO (Harris et al., 2004)) and pathway analyses using Reactome (Fabregat et al., 2016), MSigDB (Liberzon et al., 2015), and KEGG (Ogata et al., 1999)) databases.

### QUANTIFICATION AND STATISTICAL ANALYSIS

Unless indicated in the figure legend, all experiments were performed in triplicate and results are presented as mean ± standard error of mean (SEM) of absolute values or percentages of control. Each replicate was a separate tissue culture well processed in parallel. A list of all samples used for subsequent analysis can be seen in **Table S1A-B**.

Statistical P*-*values were obtained by application of the appropriate statistical tests using GraphPad Prism v.9.0. Lipidomics data was normalized to the total ion signal, glog transformed, autoscaled and analyzed with Metaboanalyst v.3.0 (Pang et al., 2021). For quality control, internal lipid standards added to each sample were used to assess technical reproducibility, achieving ≤15% relative standard deviation (RSD) across all samples. Correlation heatmaps were generated for the lipid samples using Pearson’s r. Heatmap clustering was performed using a T-test or an ANOVA, showing the most significant lipids. For lipidomics applications, Bonferroni false discovery rate (FDR) adjusted P-values lower than 0.05 and an absolute fold-change greater than or equal to 1.5 were considered significant. For RNASeq differential gene expression, gene ontology and pathway analyses, statistical significance was corrected for multiple comparisons (Bonferroni adjusted) and assessed at an α=0.05. Genes with absolute log_2_ fold changes ≥ 1 were considered significantly differentially expressed. For supernatant genome copy RT-qPCR, a two-way non-parametric ANOVA with Dunn’s post-hoc test was performed in GraphPad Prism v.6.0 and assessed at an α=0.05. Since no statistical differences were observed between technical replicates of the same condition in the supernatant genome copy RT-qPCR data, technical replicate wells were pooled in the reported analysis.

## AUTHOR INFORMATION

### Corresponding Author

Contact information for the author(s) to whom correspondence should be addressed: Rhoel R. Dinglasan and Timothy J. Garrett

## Acknowledgements

This work was funded in part by support from the University of Florida Preeminence Initiative through the College of Veterinary Medicine and the Emerging Pathogens Institute (RRD).

## Author Contributions

The manuscript was written through contributions by all authors. RRD, TG and JAL conceptualized the study. IM, HC, TH, CJS, and JA conducted the study and generated samples and data. IM, HC, TH, MM, JA, HSY, RRD and TG analyzed the data. IM, HC, TH, MM, JA, JAL, RRD and TG wrote/edited the manuscript. All authors have given approval to the final version of the manuscript.

**‡These authors contributed equally.**

