## Supplementary Figures for "Niclosamide reverses SARS-CoV-2 control of lipophagy"

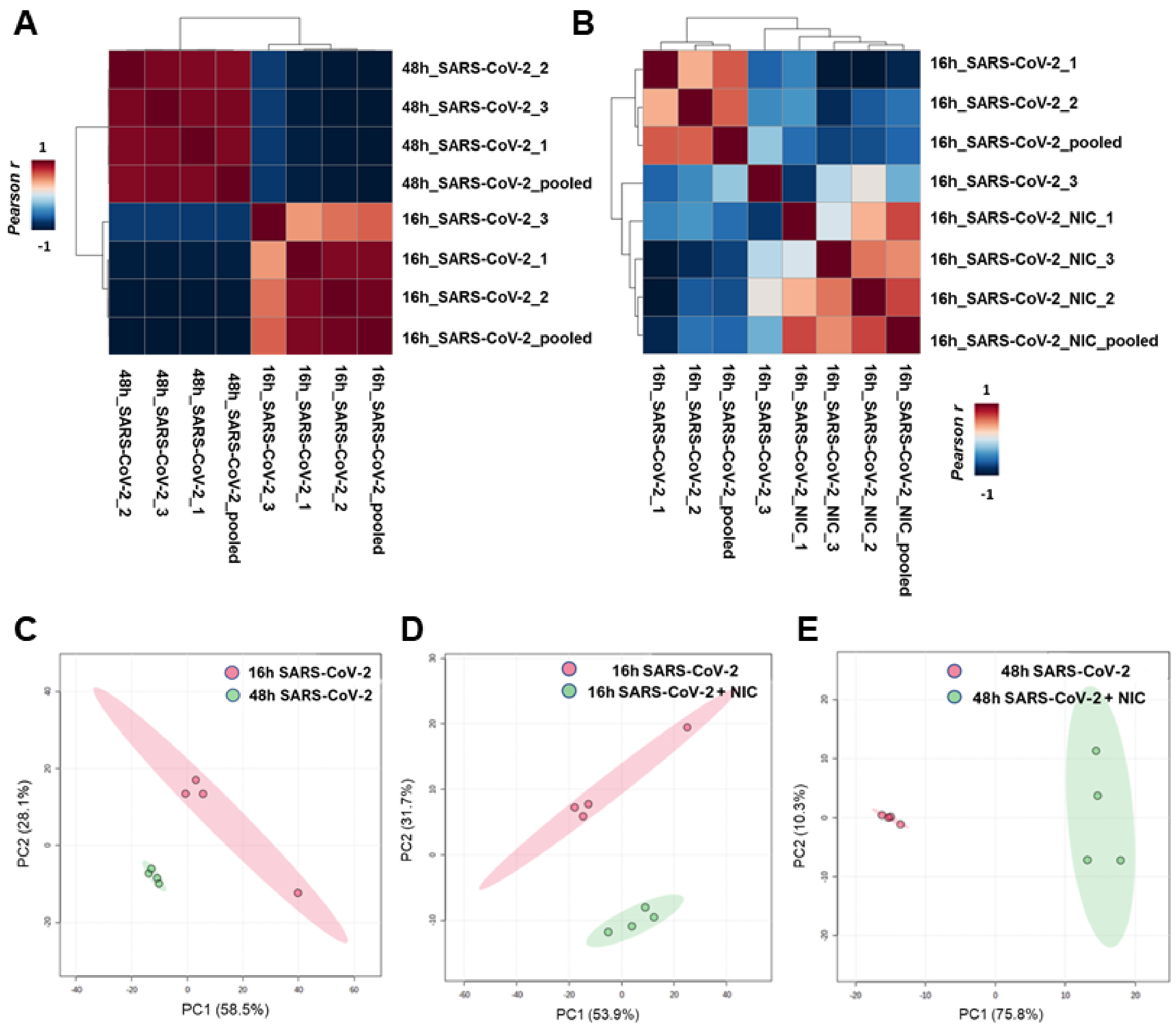


**Supplementary Figure S1. Pearson correlation analyses. (A)** Viral infection at early (16h) and late time points (48h). **(B)** Viral infection with NIC treatment at 16h. Principal components analysis (PCA) of early and late viral infection **(C)** and with NIC treatment at early (16h) and late (48h) time points **(D-E)**. Clustering was evident with viral infection and with treatment using Pearson correlation and PCA. A pooled sample from each group is identified and was used to evaluate clustering.

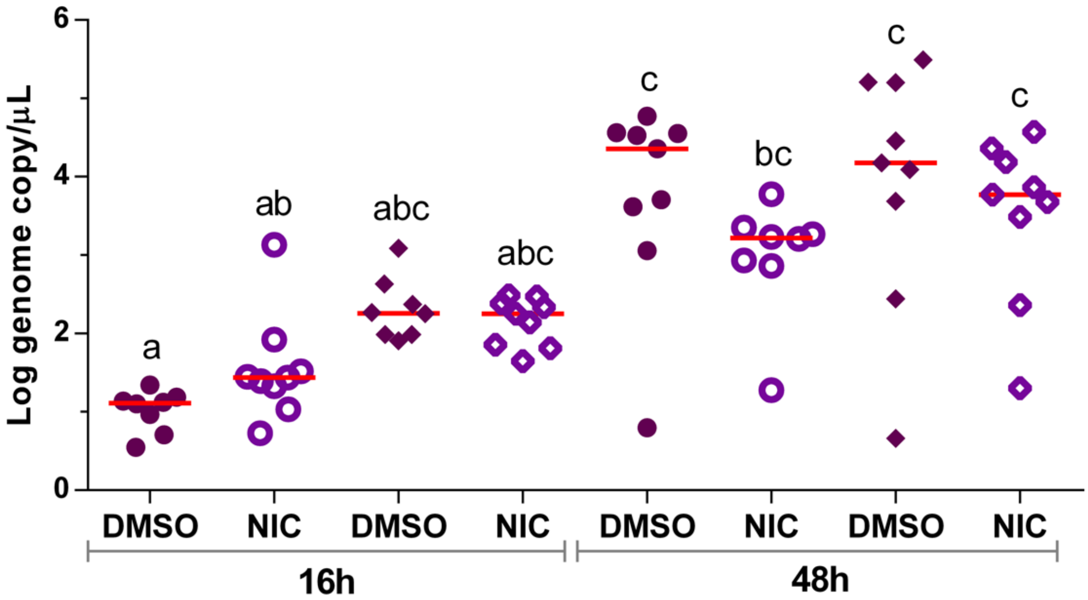


**Supplementary Figure S2. Supernatant virus genome copy in SARS-CoV-2 infected, niclosamide treated Vero E6 cell cultures.** Supernatant was collected from 8 (DMSO 16h) or 9 (all other conditions) replicate wells at the indicated timepoint after SARS-CoV-2 infection, RNA was extracted, and genome copy was quantified by RT-qPCR against the N protein normalized to a standard curve generated using a recombinant plasmid containing the N protein sequence at a known copy number. 2 technical replicates of extraction and RT-qPCR were performed on separate aliquots of supernatant from the same cultures, indicated as separate x-axis coordinates with the same condition label. 1 sample in the first replicate of the NIC 48h condition did not have any detectable genome copy and is not displayed on the graph. Different letters indicate statistically significant differences between treatments as measured by non-parametric two-way ANOVA with Dunn post-hoc test, with the threshold for significance set at $\alpha=0.05$. Red line indicates the median.

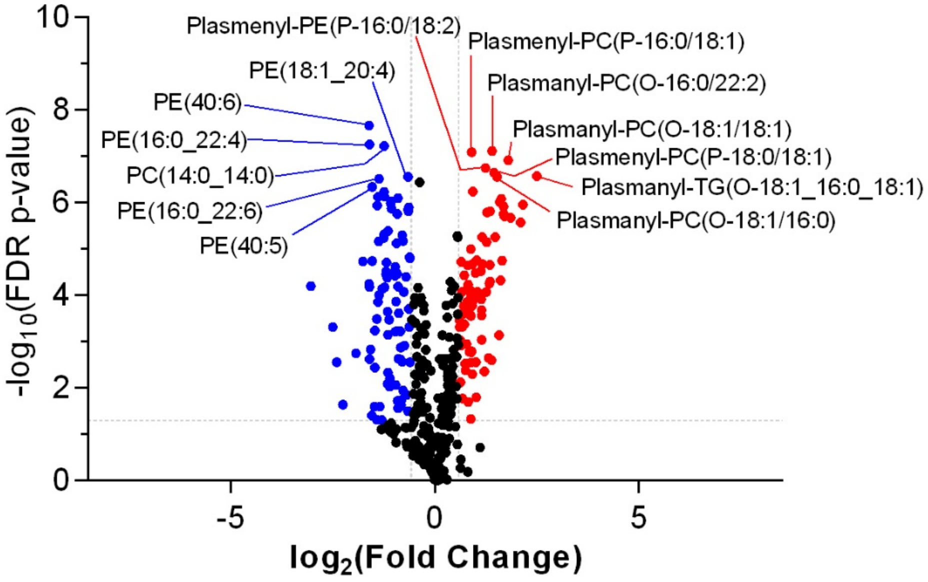


**Supplementary Figure S3. Volcano plot of 16h vs. 48h baseline lipid profile for Vero E6 in the absence of SARS-CoV-2.** A Bonferroni P-value of less than 0.05 and a fold change of 1.5 or greater were considered significant.

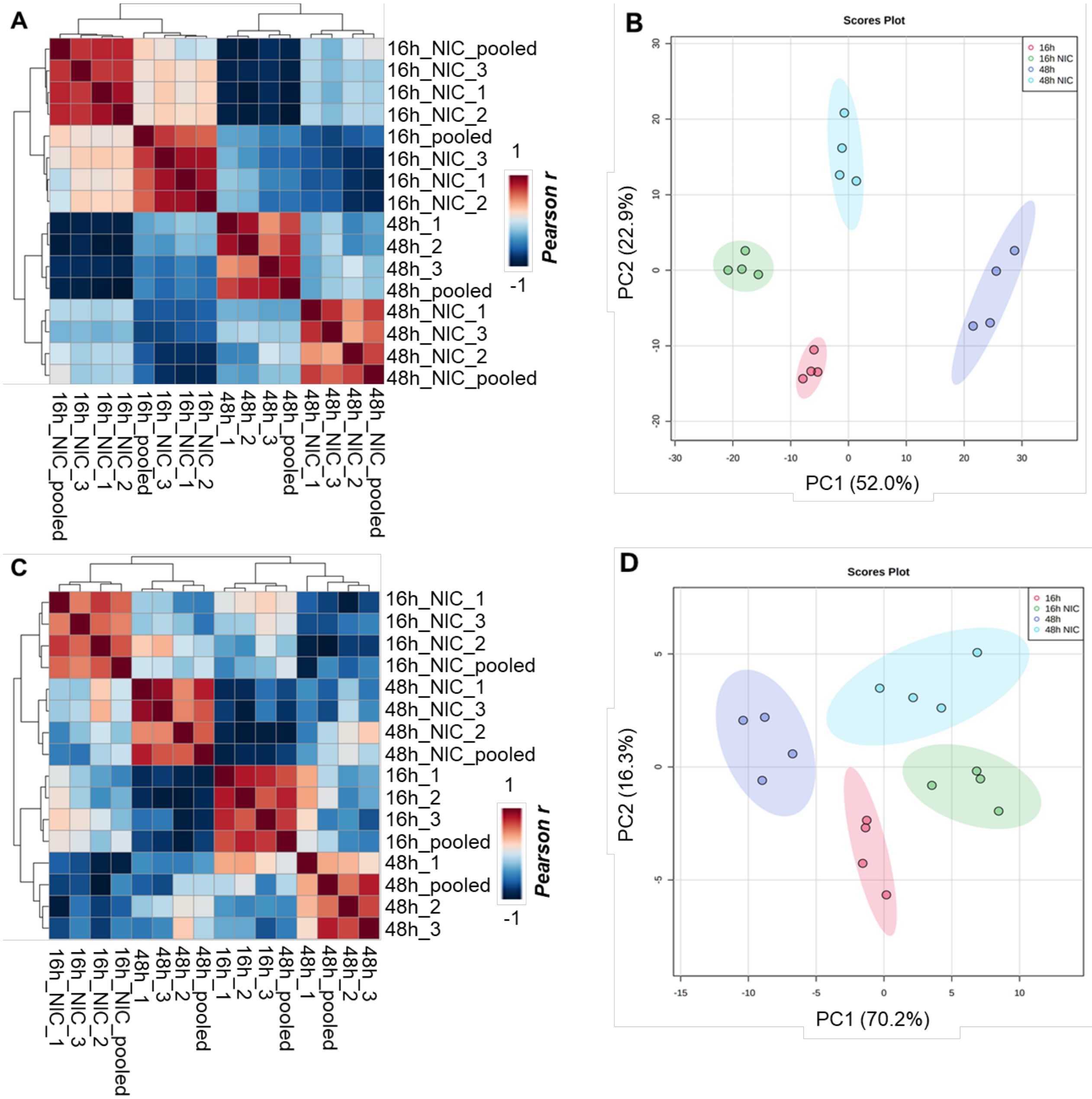


**Supplementary Figure S4. Clustering analysis of virus free Vero E6 cell cultures comparing early growth (16h) and late growth (48h) with and without NIC.** Pearson correlation and PCA including all lipids identified **(A-B)**. Pearson correlation and PCA including only TG lipids detected **(C-D)**. Clustering was evident in all cases, but correlation with all lipid expression was primarily based on time points while correlation using only TG expression was from NIC treatment.

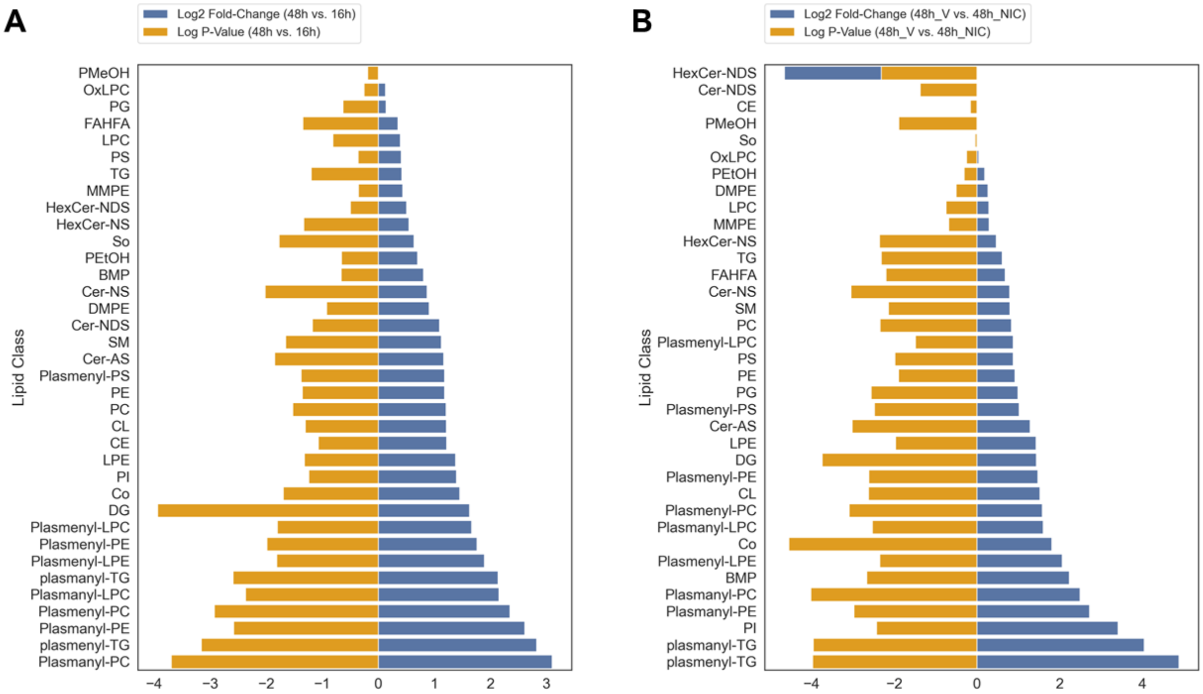


**Supplementary Figure S5. (A)** Bar graphs of the significant lipid classes at 16h vs 48h and **(B)** 48h + Vehicle (V) vs 48h + NIC. Fold change of less than 1.5 and p-value less than or equal to 0.05.

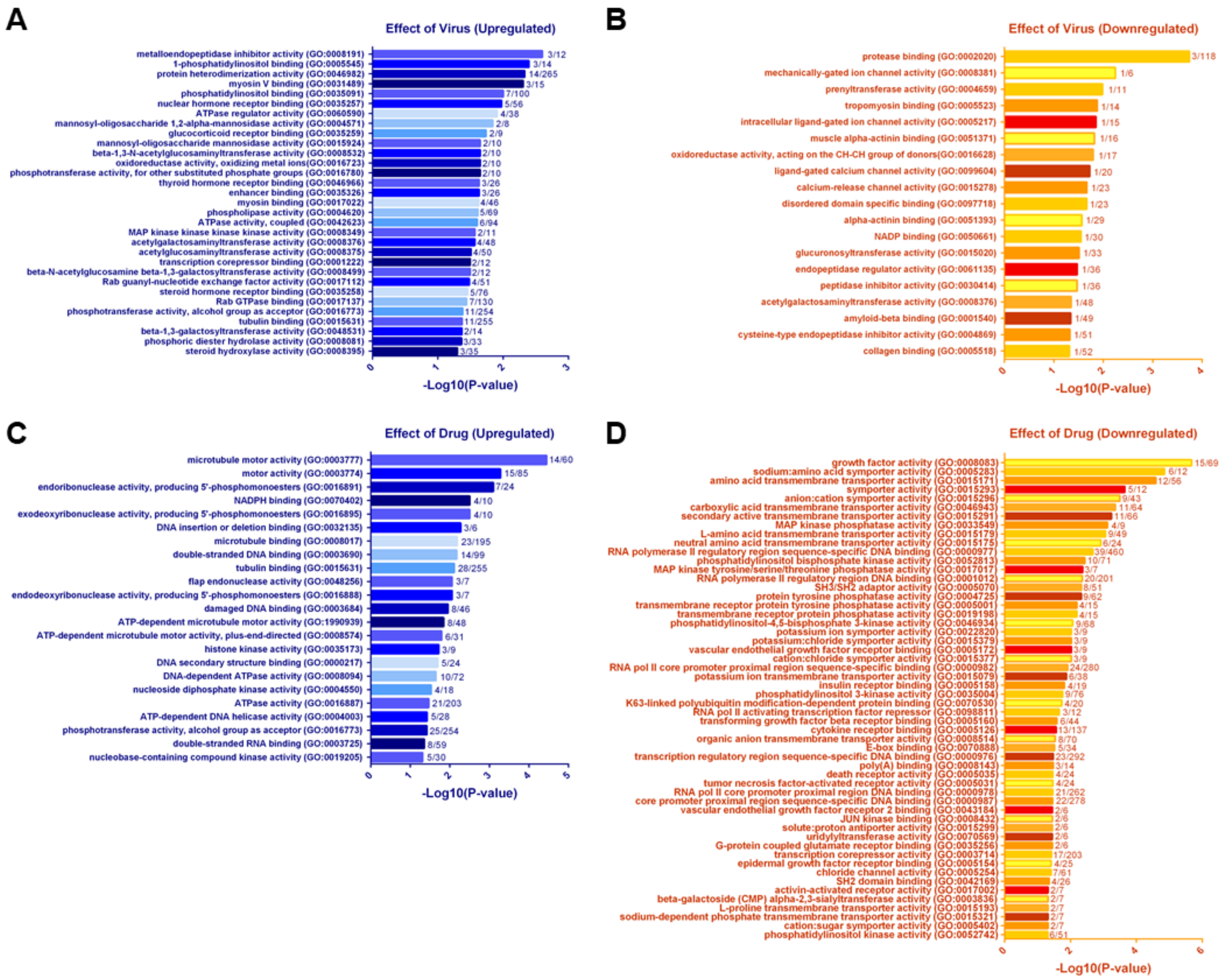


**Supplementary Figure S6. Statistically significant overrepresented gene ontology (GO) terms depicting the effect of SARS-CoV-2 infection and niclosamide treatment. (A-B)** Virus samples (with niclosamide at 16 and 48 hours, and with vehicle [DMSO] at 16 and 48 hours) were analyzed against no-virus samples and combined to analyze global virus specific transcriptome changes. Bar labels represent the number of differentially regulated genes as a function of the total number of possible genes within the listed GO term. (A) Significantly upregulated genes, colored in blue, (B) Significantly downregulated genes, colored in red and yellow. **(C-D)** Niclosamide samples (with mock at 16 and 48 hours, and virus at 16 hours) were analyzed against no-drug treated samples and combined to analyze global drug specific transcriptome changes. Bar labels represent the number of differentially regulated genes as a function of the total number of possible genes within the listed GO term. (C) Significantly upregulated genes, colored in blue, (D) Significantly downregulated genes, colored in red and yellow. Statistical significance was performed using ENRICHR and was adjusted for multiple comparisons and tested at α=0.05.
